# A novel, open source, low-cost bioreactor for load-controlled cyclic loading of tendon explants

**DOI:** 10.1101/2021.03.16.435688

**Authors:** Krishna Pedaprolu, Spencer E Szczesny

## Abstract

A major risk factor for tendinopathy is tendon overuse (i.e., fatigue loading). Fatigue loading of tendon damages the extracellular matrix and induces tissue degeneration. However, the specific mechanisms linking tendon fatigue damage with tissue degeneration are unclear. While explant models of tendon fatigue loading have been used to address this knowledge gap, they predominantly employ bioreactors that apply cyclic displacements/strains rather than loads/stresses, which are more physiologically relevant. This is because of the technical complexity and cost of building a load-controlled bioreactor, which requires multiple motors, load cells, and computationally intensive feedback loops. Here, we present a novel, low-cost, load-controlled bioreactor that applies cyclic loading to multiple tendon explants by offloading weights from a single motorized stage. Using an optional load cell, we validated that the bioreactor can effectively provide load-controlled fatigue testing of mouse and rat tendon explants while maintaining tissue viability. Furthermore, all the design files, bill of materials, and operating software are available “open source” (https://github.com/Szczesnytendon/Bioreactor) so that anyone can easily manufacture and use the bioreactor for their own research. Therefore, this novel load-controlled bioreactor will enable researchers to study the mechanisms driving fatigue-induced tendon degeneration in a more physiologically relevant and cost-effective manner.

## Introduction

Tendinopathy and tendon injuries are a significant source of pain and dysfunction. Altogether, tendon and ligament injuries account for approximately 20-30% of all musculoskeletal disorders,^8,17^ which is the leading cause of disability in the USA.^62^ Although tendinopathic pain may be manageable with non-surgical options and rehabilitation (e.g., non-steroidal anti-inflammatory drugs (NSAIDs), eccentric loading), pain recurrence is common and often chronic.^40^ Additionally, compositional and functional impairments remain after treatment, which pose a risk for future tendon tears.^6^ To improve outcomes, it is necessary to identify the underlying cause of tendon degeneration and develop new treatments to prevent disease progression and promote tissue repair.

While tendon degeneration is likely due to numerous factors (e.g., age, gender, obesity), a prominent risk factor is overuse (i.e., fatigue loading).^26^ Fatigue loading causes direct mechanical damage of the extracellular matrix^12,27,56^ and also increases the catabolic activity of tendon cells via release of matrix metalloproteinases (MMPs)^2,14,45^ and inflammatory cytokines,^14,45,50^ which further weaken the tissue. Additionally, evidence suggests that fatigue loading may be responsible for heterotopic ossification and the deposition of mucous or lipid deposits that are characteristic of tendon degeneration.^4,60,61^ However, the specific mechanisms linking fatigue loading with tendon degeneration are unclear. For example, it is still unknown whether the degenerative changes observed with tendon overuse result from a tenocyte response to elevated cyclic strains or reduced strains due to local tissue damage.^5^ To address this knowledge gap, various *in vitro* and *in vivo* models of tendon fatigue damage have been used to investigate the response of tendon cells to cyclic loading.^15,58^

*In vitro* stretching experiments of flexible substrates have produced significant knowledge regarding the response of tendon cells to cyclic loading.^52^ These studies have shown that cyclic loading of tenocytes and tendon progenitor/stem cells alters the transcriptomic and proteomic production of proteases, growth factors, matrix genes, inflammatory cytokines, apoptotic factors, and markers for non-tenogenic differentiation.^9,20,44,53,60^ Additionally, some of these studies have identified the mechanotransduction pathways responsible for these effects.^43^ However, a disadvantage with this approach is that the *in vitro* 2D conditions do not accurately represent the native tendon environment. In contrast to a dense cell monolayer, tendons are sparsely populated with cells that reside in longitudinal arrays embedded within aligned collagen fibrils and are connected via gap and adherens junctions.^21,25,33^ Additionally, tendon cells are enveloped by a pericellular matrix that physically separates the cells from the surrounding collagen fibrils.^35,39^ This indirect connection to the bulk tissue causes the local tissue strains that cells experience *in situ* to be much lower than the bulk applied tissue strains.^32,34,37^ Furthermore, cellular interactions with the pericellular matrix influences important cell signaling pathways.^10^ Therefore, while previous *in vitro* studies are valuable for understanding the response of tendon cells to direct stretching, they do not replicate the mechanical stimulation, cell-matrix interactions, and cell-cell communication that is present in the native tendon microenvironment.

At the opposite end of the spectrum, *in vivo* models maintain all the proper physiologic features of fatigue loading. *In vivo* models involving excessive treadmill running have been developed and used to demonstrate that tendon overuse leads to tendon degeneration.^49,60^ In order to more precisely control the level of mechanical loading, additional *in vivo* models have been developed that displace rat patellar tendons using a tensile device while the animal is anesthetized.^18^ These studies have provided valuable information regarding the effect of mechanical loading on tendon mechanics, the production of proteases and inflammatory cytokines, and the recovery (or lack thereof) of mechanical properties.^1,2,18,47^ However, the disadvantage of these *in vivo* models is that it is difficult to identify the mechanisms driving the degenerative response to loading. For example, in contrast to *in vitro* studies, it is not possible to measure or control the strains experienced by the cells. Furthermore, it is not possible to use small molecular inhibitors to identify the signaling pathways that transduce cyclic loading and fatigue damage into a biological response. Finally, while *in vivo* studies retain the full physiological complexity that may be involved with tendon degeneration, it is difficult to isolate the effects of mechanical loading on endogenous tendon cells from exogenous factors like circulating stem cells^48^ or immune cells.^28^

Explant models serve as an intermediate between *in vivo* and *in vitro* models and share the advantages of each approach. For example, explant models preserve the native tendon environment, including *in situ* mechanical stimuli, cell-matrix interactions, and cell-cell communication (similar to *in vivo* models), while also enabling precise control of tissue loading. Additionally, since the tissue is isolated from the body, it is possible to perform more mechanistic studies by altering culture conditions, perturbing signaling pathways via small molecule inhibitors, or performing imaging and experimental assays during testing. Finally, the more reductionist approach compared to *in vivo* models allows for the complete elimination of systemic factors or the evaluation of each factor individually through co-culture conditions. One disadvantage of explant models is that it is hard to define the culture conditions or loading protocol that truly mimics the *in vivo* condition. However, recent studies have been able to identify culture conditions that reduce the drift in mechanical properties and phenotype observed in tendon explants.^51,57,59^

Another disadvantage of explant models is that they require complicated experimental setups to accurately mimic physiological loading. *In vivo* loading of tendons is dictated by the forces imposed by physiologic activities (e.g., lifting an object). Even in cases where a part of the body must be positioned to a precise location, successful body positioning is determined by the force necessary to maintain that position and the length of the full muscle-tendon unit rather than the tendon strain. That is, *in vivo* loading of tendon is load-controlled and not strain-controlled. For this reason, experiments designed to measure the fatigue properties of tendons apply cyclic loading to a target stress rather than a target strain.^36,56^ However, most tendon explant loading systems apply cyclic displacements or strains to the samples without controlling the applied stress.^16,22,23,54^ Since tendons are viscoelastic,^19,30^ cyclically loading to the same strain leads to a decaying stress over time, which is not an accurate representation of *in vivo* fatigue loading. Additionally, by stretching all samples to the same strain, each sample experiences different stress levels due to variations in sample cross-sectional area. While some explant models apply loads/stress instead of displacements/strain,^7,13^ they are complex and expensive since they require a separate motor and load cell for each individual sample and a real-time feedback loop to drive the motors. This complication has prevented wider adoption of more physiologic load-controlled bioreactors for tendon explant studies.

In this paper, we present a novel load-controlled bioreactor that is relatively economical, easy to manufacture, does not require complicated feedback loops to operate, and fits in most incubators. Briefly, the bioreactor utilizes hanging weights that are cyclically offloaded by a single motorized table to load the explants to a consistent and sample-specific stress. This setup allows precise cyclic loading of tendon explants with a single motor and even without the need for a load cell. Additional displacement sensors can be used to record the resultant displacement of each sample, which enables the calculation of strain and modulus in each cycle over time. We validated that the bioreactor can effectively provide load-controlled fatigue testing of tendon explants and is able to keep the tendon explants viable for ^~^26 hours of loading. We have also provided open access to the CAD files, engineering drawings, bill of materials, and operating software (https://github.com/Szczesnytendon/Bioreactor) so that anyone can easily manufacture and use the bioreactor for their own research.

## Materials and Methods

### Bioreactor Design

The full assembly of the bioreactor and each component of the loading mechanism are shown in Figure 1. Each loading mechanism consists of a pair of autoclavable stainless steel grips that hold the tendon submersed in culture medium within a rectangular well plate. These grips are mounted on linear bearings (RX series, Tusk Direct) to support off-axis loads and moments while providing smooth linear translation and force transmission. One of the linear bearings is fixed to a rigid aluminum backplate either directly or via a load cell. The other linear bearing is free to move and is connected to a hanging weight by a fishing line that passes over a low friction pulley. There are four such loading mechanisms in the bioreactor so that four tendons can be cyclically loaded simultaneously. Cyclic loads are applied by offloading the hanging weights via a motorized stage (VSR40A-T3, Zaber) that moves vertically with a sinusoidal displacement profile controlled by LabVIEW. Static loads can be applied by hanging weights that do not rest on the stage. Note that because the hanging weights provide a known cyclic load to each sample, the use of a load cell is optional and only necessary to investigate potential frictional losses in the load path. Additionally, the mobile grip is connected to a linear variable differential transformer (LVDT) (HR 200, TE connectivity) that is mounted to the bioreactor base. This enables real-time measurement of the sample displacements and, therefore, calculation of the tissue strain and modulus during each loading cycle. A video of the bioreactor in operation is available in the Supplementary Materials. Additionally, the computer-aided design (CAD) models, engineering drawings for each component, and the complete bill of materials are available at https://github.com/Szczesnytendon/Bioreactor.

**Fig 1.**
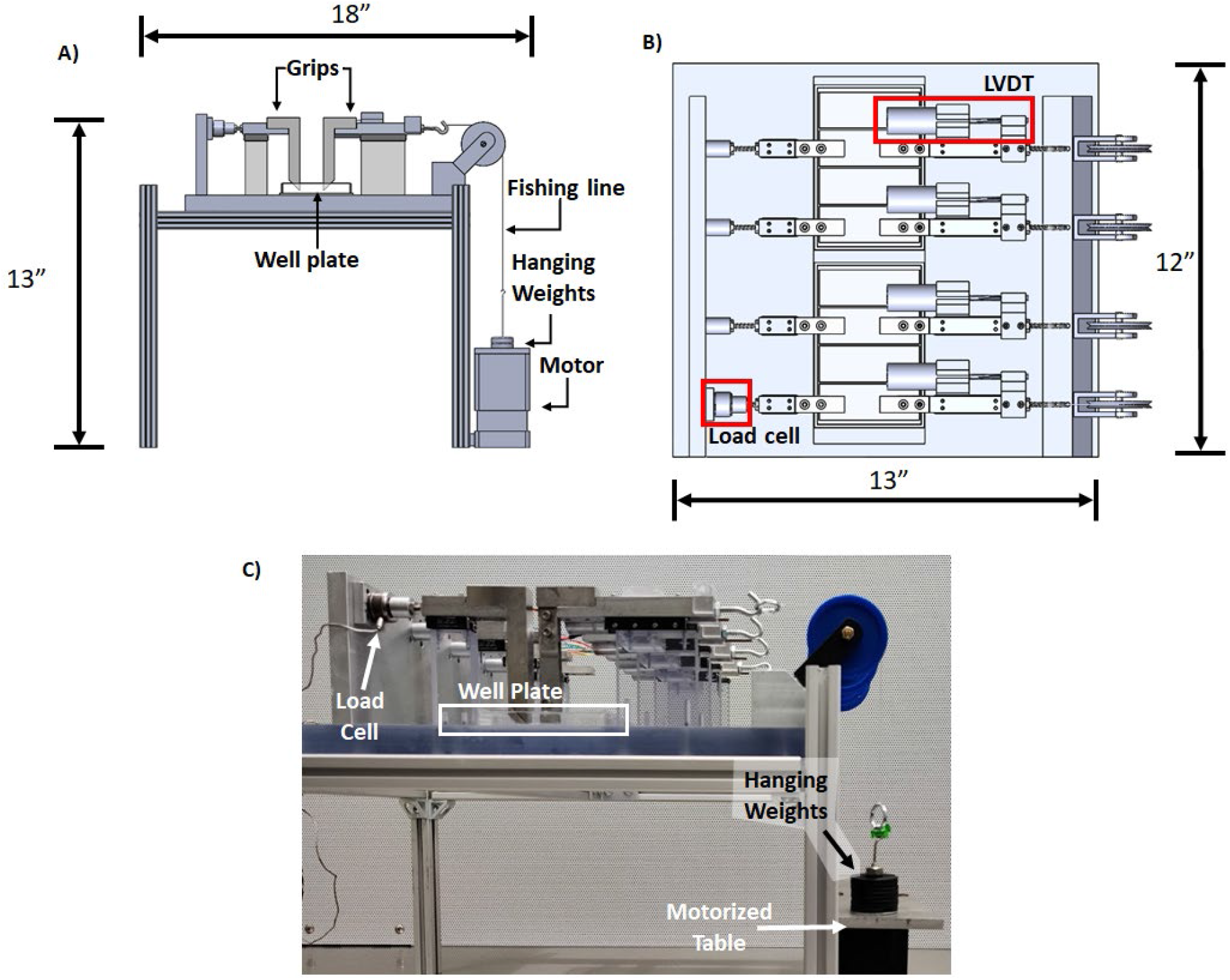
Design of the bioreactor.

(A) Side view of the bioreactor and its parts. (B) Top view of the bioreactor. (C) Image of the fully assembled bioreactor.

### Gripping Techniques

Tendons can be tested within the bioreactor with either the bony attachment intact or removed using two different gripping techniques (Fig. 2). For tendons with a bony attachment, the piece of bone is secured by grips composed of two parallel plates with vertical serrations (Fig. 2A, 2C). To secure tendons with no bony attachments or by the myotendinous aponeurosis, the tissue is gripped by an angled stainless-steel clamp lined with sandpaper (Fig. 2B, 2C). To adhere the sandpaper to the clamps, double-sided tape (467 MP, 3M) is attached to the sandpaper and then the sandpaper is cut using a laser cutter into pieces that match the shape of the clamps. Finally, the cut sandpaper pieces are sterilized by soaking them in 70% ethanol for 30 minutes and air dried in sterile conditions.

**Fig. 2:**
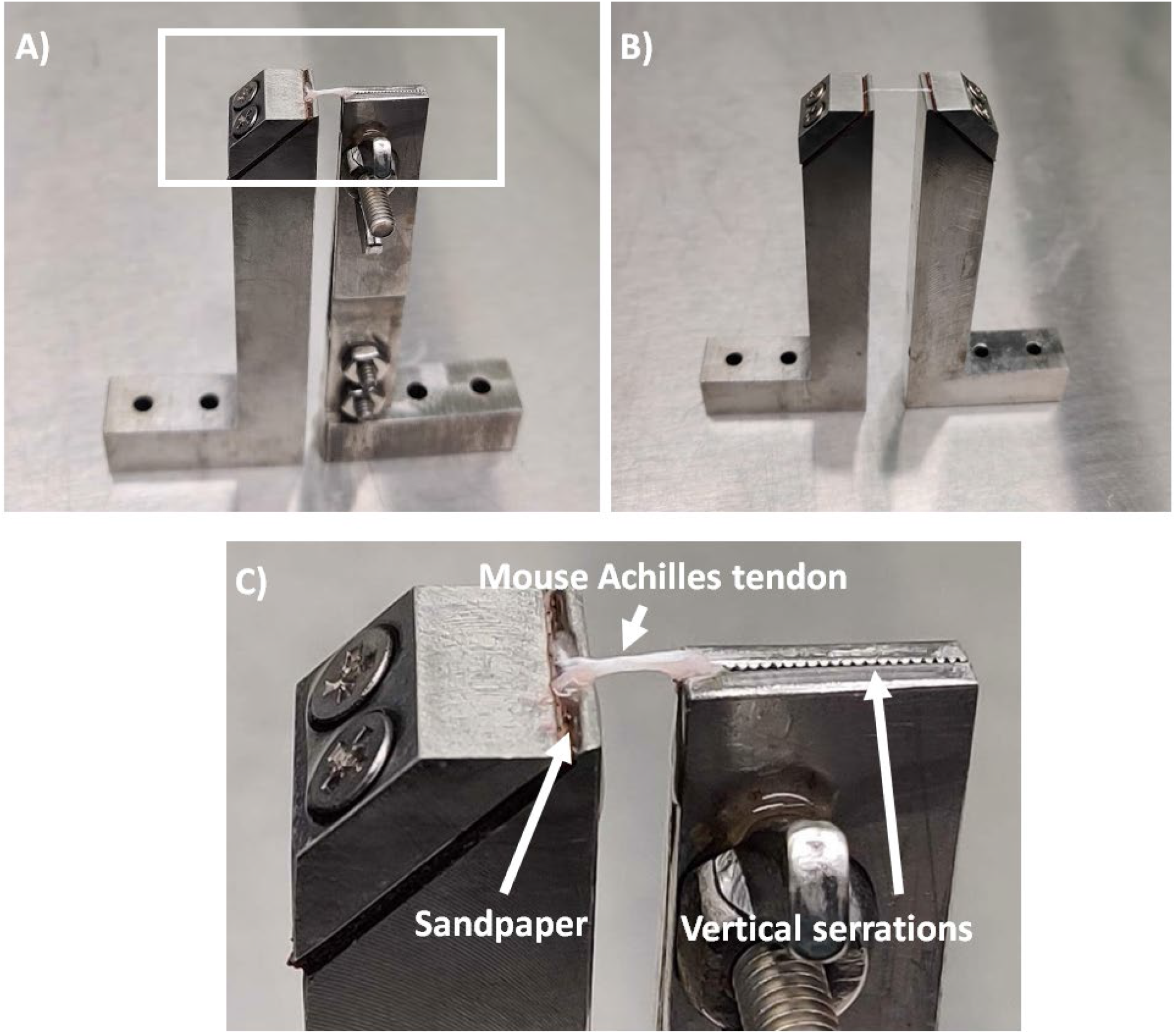
Gripping techniques.

(A) Grips used to secure tendons with an intact bony attachment or (B) with the bony attachment removed. (C) Zoomed in image of white box in (A) showing the details of both grips. Note the tendons depicted in the images are a (A) mouse Achilles tendon and (B) a rat tail tendon fascicle.

### User Interface Software

A LabVIEW code (also provided on the GitHub link) is used to run the bioreactor, display the results, and record the data (Fig. 3). The output from the load cell conditioner (GM, Honeywell) and the LVDT conditioners (LVM110, TE Connectivity) are connected to a data acquisition (DAQ) board (USB6000, National Instruments). The data from the DAQ board is processed by the LabVIEW code to display and record the position of the free moving grips and the loads experienced by the tendon. Further, the LabVIEW code also controls, records, and displays the amplitude and the frequency of the sinusoidal displacement of the motor. The data from the motors, the load cell, and LVDTs are recorded and saved as a .lvm file.

**Fig. 3:**
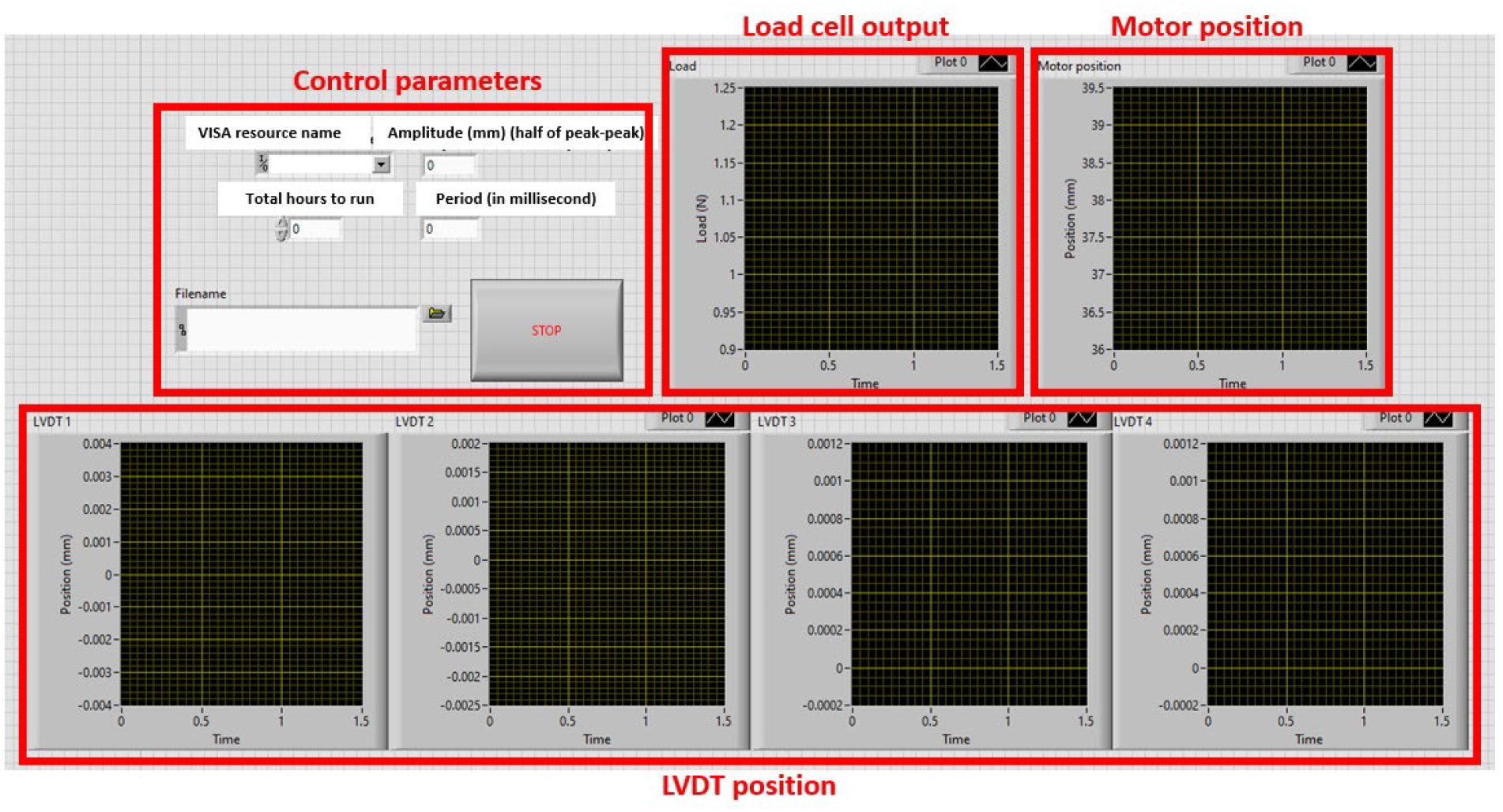
User interface of the LabVIEW code.

The amplitude and frequency of the motor displacement can be specified on the front panel. Additionally, the outputs of the load cell, motor, and each LVDT are displayed in real-time.

### Cyclic Fatigue Loading of Mouse Achilles Tendons

Fatigue loading tests of tendon were performed to demonstrate the ability of the bioreactor to apply consistent cyclic loads, accurately measure the data, and maintain tissue viability. Twenty-four mouse Achilles tendons were extracted from twelve male C57BL/6 mice (age: 8 ± 4 months) under an approved IACUC study. The calcaneus was retained and most of the gastrocnemius and soleus muscles were scrapped off using a scalpel blade to make the myotendinous aponeurosis easy to grip. The cross-sectional areas of the tendons were determined by a non-contact laser transducer (LJ-V7060, Keyence). The calcaneus and myotendinous aponeurosis were fixed in the serrated grips and sandpaper grips, respectively. Before placing the tendons in the bioreactor, the gauge length was determined by measuring the distance between the calcaneus insertion and the lip of the stainless-steel clamp visually with a vernier caliper while ensuring there was no slack in the tissue. All the tests were performed in culture medium (low glucose Dulbecco’s Modified Eagle Medium, 1% penicillin-streptomycin, 200 mM Glutamax, 25 mM HEPES, 1 mM magnesium L-ascorbate) at 30°C and 5% CO_2_. The bioreactor applied a cyclical load of 2 MPa at 0.5 Hz for 47,466 cycles (94,932 seconds) to twelve samples. Based on preliminary experiments, this number of cycles corresponds to 70% of the fatigue lifetime for mouse Achilles tendons under these loading conditions. Prior to loading eight of the twelve samples, the bioreactor was kept in the incubator for 3 hours to allow for thermal equilibration of the system and prevent drift in the displacement data due to thermal expansion of the LVDT components. Four of the twelve samples ruptured before the loading ended. The remaining eight samples were analyzed for mechanical data. Viability was calculated for six of the eight non ruptured samples. An additional twelve samples were used as controls, with six samples statically loaded to approximately 0.06 MPa and six samples immersed in culture media outside the bioreactor for the same duration as the fatigue loaded samples.

### Data Analysis

The tissue displacement was calculated as the difference in the LVDT position during testing and the initial position. The tissue strain was then calculated by dividing the displacement by the initial gauge length of each tendon. The peak strain was defined as the maximum strain value measured during each cycle of the motor displacement. Finally, the tissue modulus was determined for each cycle by dividing the maximum applied stress (2 MPa) by the difference between the maximum and minimum strain value recorded during that cycle. Plots of the peak strain and tissue modulus versus time were smoothed using a moving average of 50 and 1000 data points, respectively.

### Validation of Load Applied to Tendon

The load cell data were used to validate that the magnitude and frequency of the load transmitted to the tendon samples were accurate. Specifically, four freshly frozen rat flexor carpi ulnaris (FCU) tendons harvested under the same approved IACUC protocol were cyclically loaded for 1000 cycles with a 1 N weight and a motor frequency of 0.5 Hz. The maximum load was determined by averaging the values within the peak of each loading cycle. The duty cycle of the loading profile was calculated as the amount of time per cycle that the load was above 50% of the maximum load. Finally, the load output was averaged over the duration of loading and compared to the time-averaged force magnitude of an ideal sinusoidal waveform (i.e., 0.5 N).

### Tissue Viability

Following loading, the mouse Achilles tendon samples were stained with 0.2% of 4 mg/ml fluorescein diacetate (FDA) (596-09-8, ACROS Organics) and 0.5% of 1 mg/ml propidium iodide (PI) (P3566, ThermoFisher Scientific) added to the media in the bioreactor to label live and dead cells, respectively. After 10 mins of staining, the gripped tendons were transferred to a tensile testing device^31^ mounted on a confocal microscope (A1R HD, Nikon Instruments). A minimal load was applied to the gripped tendons to remove any slack before imaging. Free floating samples were held under a coverslip with small weights and then imaged. Three-dimensional images (2.989 μm z-step, 2.49 um/px) were captured on the confocal microscope, and the images were thresholded to eliminate background and identity regions of positive staining. Tissue viability was calculated as the ratio of FDA-positive pixels to the total fluorescence (i.e., the sum of pixels positive for FDA and PI). The region between the grips was used for calculating viability in gripped samples and the mid-substance was analyzed in the free-floating samples. Thresholding and region selection was done by two blinded observers. Viability was calculated as the average of the values obtained by both observers.

### Statistics

A one sample t-test was used to determine if the difference between the time-averaged force measurement and that of a matching sinusoidal waveform was significantly different than zero. Viability of the statically loaded and fatigue loaded samples was compared to free floating samples using a Brown-Forsythe and Welch ANOVA test with a Dunnett’s multiple comparisons test. Statistical significance was set as p < 0.05.

## Results

### Tendon Fatigue Loading

Figure 4 shows the raw data output during loading of a representative mouse Achilles tendon sample. The motor exhibited a smooth sinusoidal displacement with an amplitude of ±4 mm and a frequency of 0.5 Hz (Fig. 4A). The LVDT position progressively increased during loading and exhibited a square waveform (Fig. 4B-C). The load applied to the tendon (as measured by the load cell) also exhibited a square waveform (Fig. 4D-E). While the load cell output changed over time, pilot testing at room temperature suggests that this was a result of the system reaching thermal equilibrium and was not an actual change in load (Fig. S1).

**Fig. 4:**
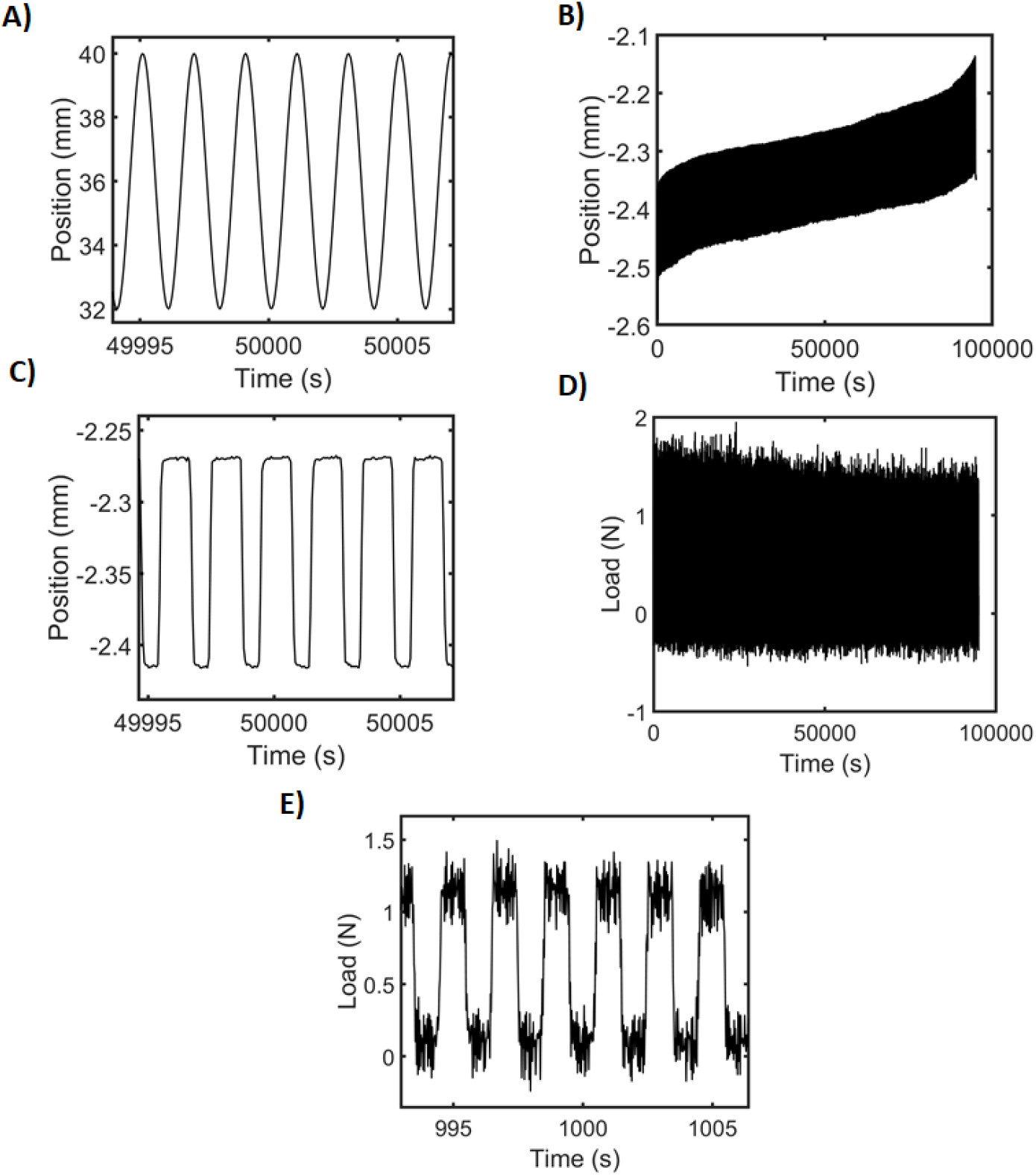
Raw data output from bioreactor.

Representative graph of the (A) motor position, (B) LVDT position with (C) detail of LVDT waveform, and (D) load cell reading with (E) detail of load waveform of an Achilles tendon sample loaded to 2 MPa.

Similar to the raw displacement data, the peak strain of each cyclically loaded sample progressively increased over time. There was an initial rapid increase in peak strain (primary phase) followed by a slow linear increase (secondary phase) (Fig. 5A). Two samples exhibited a rapid increase in strain at high cycle numbers (tertiary phase) while four samples ruptured during testing and were not analyzed. Additionally, one sample slipped during loading and was excluded. While the tissue modulus exhibited a variable trend for each sample, overall, the value was largely unchanged (Fig. 5B). Finally, the samples that were not allowed to thermally equilibrate prior to testing (highlighted in red) exhibited a drift in displacement during the early cycles (asterisks).

**Fig. 5:**
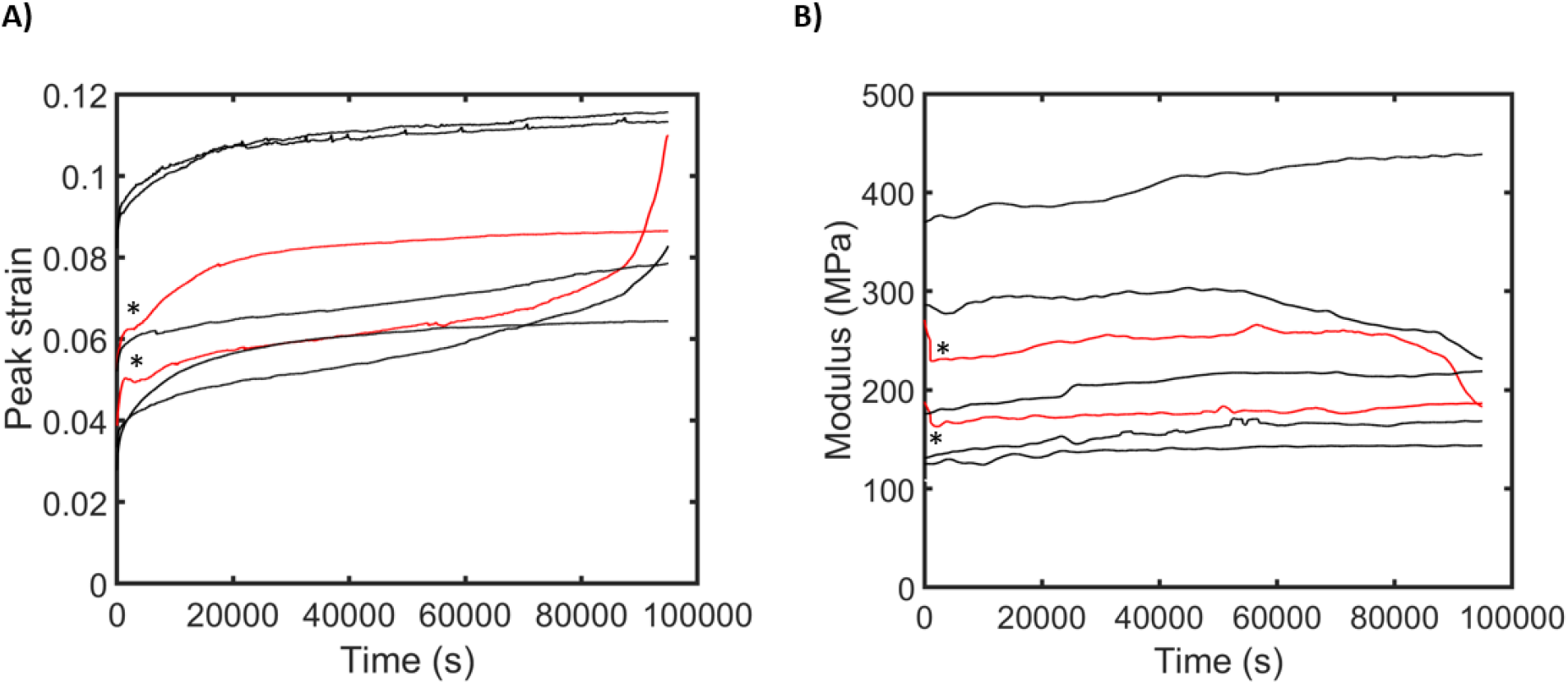
Mechanical properties of mouse Achilles tendons during fatigue loading.

(A) Peak strain versus time for each sample. (B) Tissue modulus versus time for each sample. Red lines indicate the samples that were loaded prior to the system reaching thermal equilibrium. Asterisks point out the drift in displacements during early cycles.

### Validation of Load Applied to Tendon

Similar to the testing of the mouse Achilles tendons, the load applied to the rat FCU tendons exhibited a square waveform (Fig. 6). The average maximum load recorded was 0.96 ± 0.01 N for a hanging weight of 1 N, and the duty cycle was 0.39 ± 0.05. The time-averaged load magnitude was 0.39 ± 0.05 N, which was significantly different from the mean load (0.5 N) of an ideal sinusoidal loading profile (p < 0.05).

**Fig. 6:**
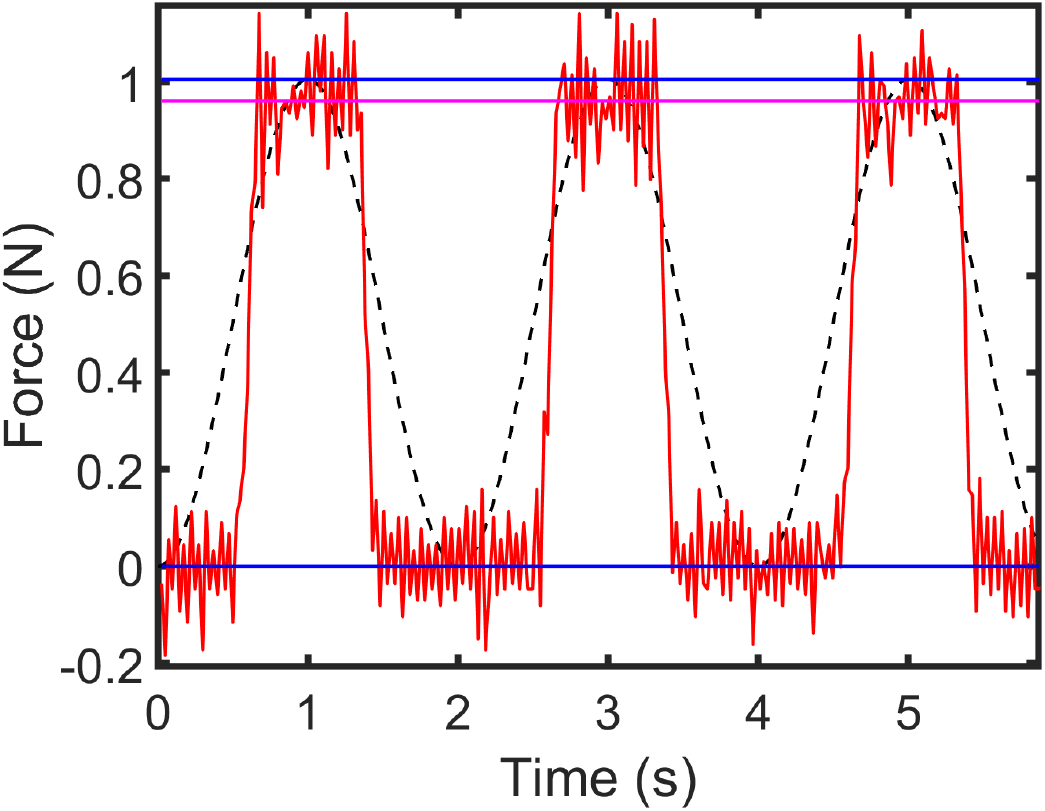
Validation of applied load.

Representative graph of the applied loading profile (red) and the ideal sinusoidal loading profile (black). Blue lines represent the minimum (0 N) and maximum load (1 N) of the ideal sinusoidal profile. The magenta line represents the average maximum load (0.96 ± 0.01 N) recorded by the load cell.

### Tissue Viability

The samples remained viable following fatigue loading (Fig. 7). There was no significant difference between the viability of fatigue loaded (81 ± 10 %) and static samples (78 ± 18 %) when compared to free floating samples (89 ± 5 %) (p = 0.18 and 0.31, respectively).

**Fig. 7:**
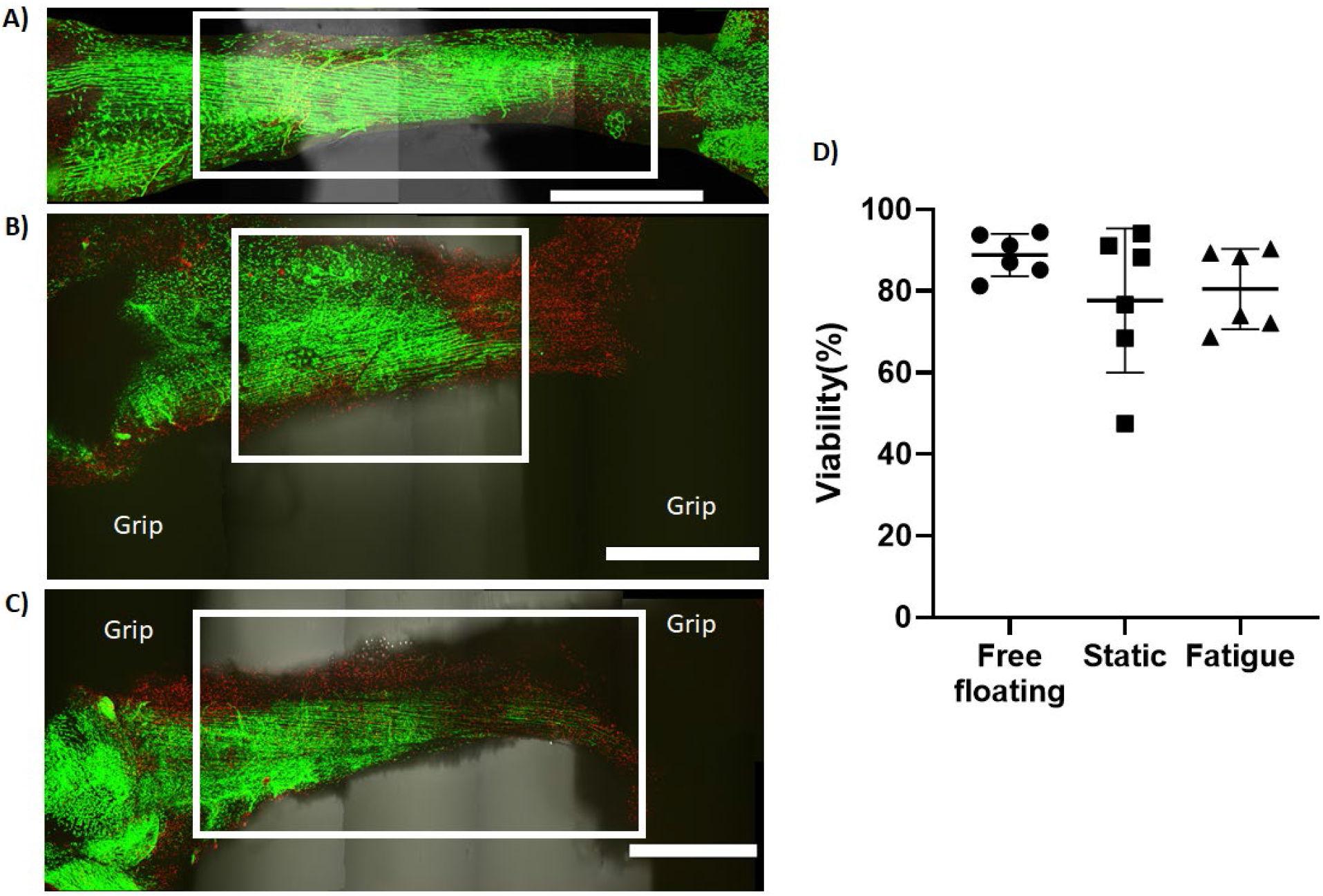
Viability of tendons in the bioreactor.

Representative image of FDA/live (green) and PI/dead (red) staining of a (A) free-floating tendon, (B) statically loaded tendon, and (C) fatigue loaded tendon. The white box indicates the region of the image that was used for calculating the viability of static and fatigue samples, which corresponds to the region between the grips. (D) There was no significant difference between the viability of the static and fatigue loaded samples compared to the free-floating samples. Scale bar = 1 mm

## Discussion

In this paper, we presented the design and validation of a simple load-controlled tensile bioreactor for studying the effect of cyclic loading on tendon explants. Specifically, the bioreactor was capable of fatigue loading mouse Achilles tendon explants and measuring their mechanical response in real-time. Consistent with prior data, the tendons exhibited a typical triphasic fatigue response^42^ and modulus values in the range of 200-400 MPa^3,11^ (Fig. 5). Importantly, the bioreactor was also able to maintain tissue viability during loading (Fig. 7). Finally, due to minimal friction in the assembly, 96% of the applied load was transmitted to the sample. Therefore, this bioreactor can effectively provide load-controlled cyclic testing of tissue explants that is more physiologic compared to displacement-controlled systems.^23,29,50,54,55^ In the future, we will use this bioreactor to cyclically load live tendon explants to understand how fatigue damage induces tendon degeneration and leads to tendinopathy.

The simplicity of the bioreactor is its primary innovative design feature. Unlike other load-controlled systems,^7,13^ this bioreactor only requires one motor and no-load cells or real-time feedback control. Stress levels are simply adjusted by calibrating the hanging weight to the sample cross-sectional area. A minimum non-zero stress value can be applied by adding an additional weight on a shorter string that is not offloaded by the motor when it moves. This makes the bioreactor effective and versatile while also inexpensive and easy to use. Furthermore, we have made the CAD models, engineering drawings, bill of materials, and LABVIEW software freely available at https://github.com/Szczesnytendon/Bioreactor. Therefore, all the necessary information is provided “open-source” for any user to build, operate, and modify the bioreactor for their own personal interests.

One limitation to the bioreactor is that the loading profile is a square wave rather than a true sinusoidal function. Additionally, the time-average load applied to the sample is 80% of what a sinusoidal waveform would produce. This is because the weights spend more time offloaded by the motor than hanging in the air, thereby reducing the effective duty cycle of the loading profile to approximately 40%. This would have the effect of delaying the number of cycles necessary to induce sample failure for a given applied stress. Therefore, if strict waveform control is important for the experiment, then a setup with a feedback control system would be a better choice. Due to the nature of loading, another limitation is that the loading frequency is limited by the bounce/recoil of the tendon as the weight comes off the motor at high speeds. However, based on our experience, this was not an issue up to 1 Hz, which is a typical loading frequency for tendon fatigue loading.^24,41,45–47,50^ Finally, the open bath design can lead to significant evaporation of the culture medium over time. This can be mitigated by the use of an incubator with active humidity control. Alternatively, the bioreactor can be redesigned with a closed bath^38,54,59^ while still using the same hanging weight principle. Despite these limitations, this simple load-controlled bioreactor can enable accurate load-controlled testing of tissue explants with significantly less cost and complexity compared to existing systems.

## Supporting information

Supplemental video of the bioreactor

## Acknowledgements

This work was supported by the Alliance for Regenerative Rehabilitation Research and Training.

## Supplementary information

**Fig. S1:**
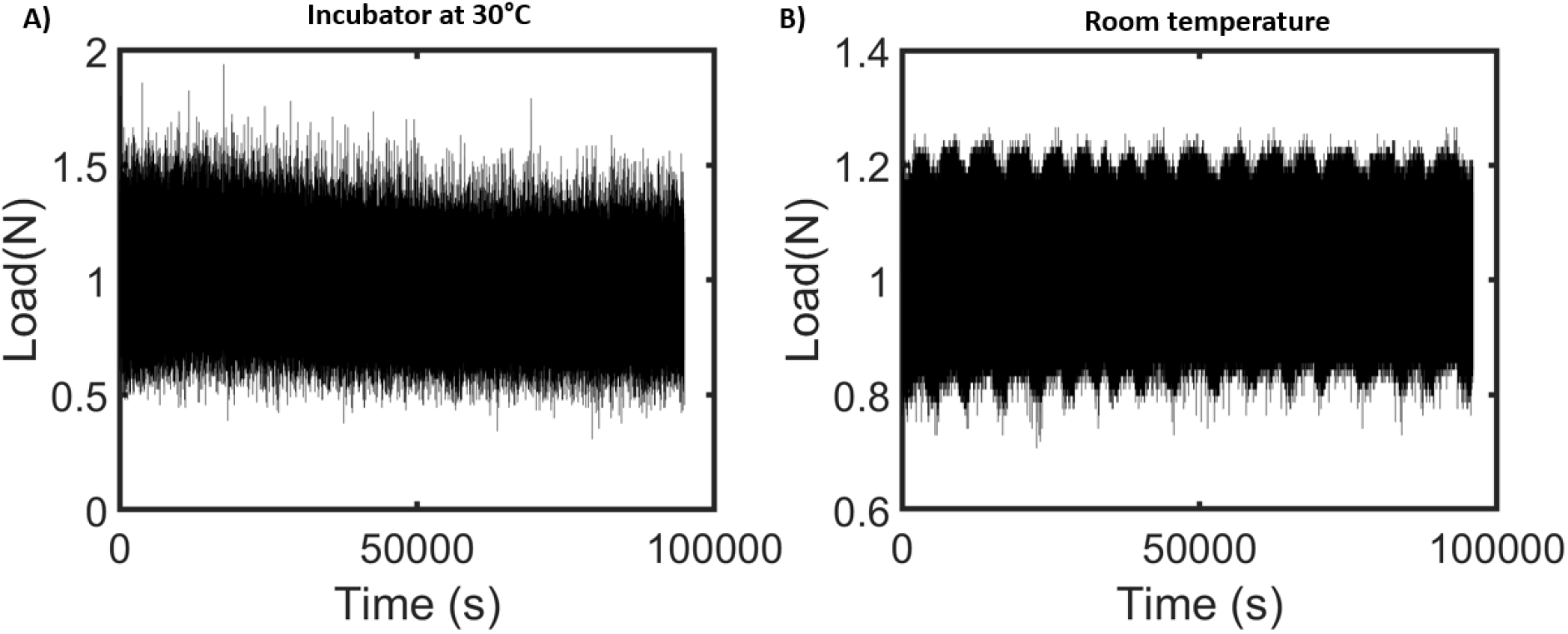
Effect of the system reaching thermal equilibrium on the load cell readout.

**A** 1 N load was applied to the load cell A) in an incubator set at 30°C and B) at room temperature. A small drift in the load cell output was observed for the load cell in the incubator, whereas the load cell readout was unaffected at room temperature.

